# Bit-twiddling on Nucleotides

**DOI:** 10.1101/082214

**Authors:** Fabian Klötzl

## Abstract

Bits, nucleotides and speed.

## 1 Introduction

Analyzing sequence data is one of the main focuses of bioinformatics. In this note we shall analyze four common tasks: computing the GC-content, hashing k-mers, computing the reverse complement and counting transversions. We present improved algorithms that show superior performance compared to na¨ıve approaches.

The sources of this paper including LATEX and program code are available at https://github.com/kloetzl/biotwiddle.

## 2 Materials and Methods

A lot of bioinformatics tools work with nucleotide data. Usually, each nucleotide is represented by one character in IUBUB nomenclature. For the four standard DNA bases these are `A, C, G`, and `T`. These letters are commonly encoded in ASCII and take up one byte of memory. Assuming a C memory model, a sequence of nucleotides is stored as a string, continuous memory, with each byte representing one character delimited by a `NUL`-byte.

### 2.1 Definitions

An alphabet is a non-empty set of characters, commonly written as Σ. A finite sequence of characters *w* = *w*_1_*w*_2_ … *w*_n_ with *w*_i_ ∈ Σ is called a string with length *n* = |*w*|. In this note the alphabet is assumed to consist of the four nucleotides from DNA: Σ = {`A, C, G, T`}.

Each character is also the representative for a seven bit sequence, namely their ASCII encoding. We allow the execution of bitwise logical operators on these sequences. For example, let *c* and *d* be two bit sequences. Then *c* & *d* is the result of a bitwise-logical-and.

### 2.2 GC-content

The GC-content of DNA is the proportion of guanin and cytosin among all nucleotides. Let *w* be a DNA sequence of length *n.* Furthermore, *S*: Σ →{0, 1} maps a nucleotide 1 iff it is a `C` or `G`. So the GC-content of the sequence *w* is 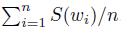.

This definition easily translates into code. For every character in the given string, check if it is `C` or `G`; if so, increase a counter. Once all characters are processed, compute the final ratio.

~~~
**double** *gc***(const char** *\*seq*) {
    **size_t** *gc* = 0;
    **const char** *\*ptr = seq;*

    **for (;** *\*ptr; ptr*++) {
        **if** *(*ptr* == ’G’ **||** *\*ptr* ***==*** ’C’) **{**
            gc++;
        }
    }

    **return (double)***gc* **/** *(ptr – seq);*
}
~~~

This looks like three comparisons are made against each character, one for `NUL` and two against `C` and `G`. However, compilers can optimize them into just two comparisons. Luckily, in ASCII, `C` and `G` differ by only one bit. This enables optimizing compilers (or us) to rewrite the comparisons to use a bit mask. We ignore the bit which differs between `C` and `G` and check if the remaining bits equal the common bit pattern.

~~~
**double** *gc***(const char** *\*seq)* {
    **size_t** *gc* = 0;
    **const char** *\*ptr* ***=*** *seq;*

    **for** (; *\*ptr; ptr*++) {
        **char** *masked = *ptr* & ’G’ & ’C;
        **if** (*masked ==* (’G’ & ’C’)) {
            *gc*++;
        }
    }

    **return (double)***gc* / *(ptr – seq);*
}
~~~

As most sequence data is encoded as ASCII, computing the GC-content the new way may result in great performance benefits for a whole lot of bioinformatics applications.

### 2.3 Hashing

A common procedure on k-mers is to hash them. This allows for compact representation in memory, or can be used as an index into hash-based data structures. As the DNA-alphabet consists of only four characters, two bits suffice to represent one nucleotide. With this we can define the following mapping for *k* ≥ |*w*|.

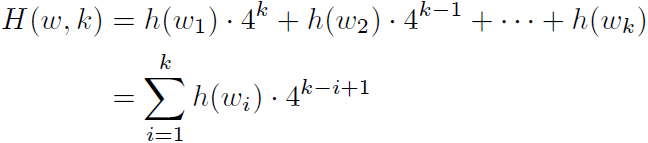

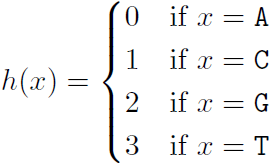

Again, this definition easily translates into code: Iterate over the first *k* characters of a string, compute the map, and combine them into one number.

~~~
**size_t** *hash(***const char** *\*seq*, **size_t** *k*) {
    **size_t** *return_value* = 0;

    **while** (*k*––) {
        **char** c = *\*seq++*
        **size_t** *val* = 0;
        **switch** (c) {
            **case** ’A’: *val* = 0; **break**;
            **case** ’C’: *val* = 1; **break**;
            **case** ’G’: *val* = 2; **break**;
            **case** ’T’: *val* = 3; **break**;
    }
        *return_value* << = 2;
        *return_value* |= *val;*
    }

    **return** *return_value;*
    }
~~~

The **switch**-statement is convenient for humans to read, but not the most compact way to achieve the desired mapping. Instead of essentially doing four comparisons, we can resort to bit-twiddling, giving us the same mapping with fewer machine instructions.

Remember, the nucleotides are stored as ASCII characters in memory. The actual values are shown in the table above. In the table, it can be seen that the lower bits 2 and 3 of each character uniquely identify it [4]. Further more, they almost achieve the desired mapping given as *h(x)* above. However, if bit 3 is set (i.e. `G` or `T`) the bit 2 is wrong. Thus it needs to be flipped conditionally using a bitwise-exclusive-or as shown in the following code.

~~~
**Size_t** *hash*(**const char** \**seq*, **size t** *k*) {
    **size_t** *return_value* = 0;

**while** (*k*––) {
        **char** *c* = \**seq*++;
        *c* &= 6;
        *c* ^= *c* ≫ 1;
        **size_t** *val* = *c* ≫ 1;
        *return_value* ≪= 2;
        *return_value* |= *val*;
    }

    **return** *return value*;
}
~~~

**Table 1.**
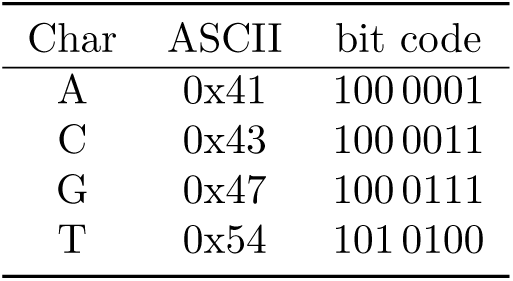
ASCII Table (excerpt)

Finally, we have achieved the mapping we set out for. In C, this code compiles down to just five bit-twiddling instructions, making it the fastest order-preserving method available. Furthermore, unlike the switch-statement, this code can be vectorized.

### 2.4 Reverse Complement

DNA consists of a forward and a reverse complement strain. For this note we use the following, technical, definition. Let *w* ∈ Σ* be a DNA sequence with 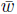 being its reverse complement, given

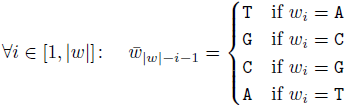

A simple implementation for the reverse complement follows a similar scheme as the hashing procedure above. Iterate over all characters in the forward sequence, map everyone to its complement, and write the result to the reverse string.

This time we focus our efforts on the complementing. Specifically, complementing a nucleotide can be compactly written as `A ↔ T` and `C ↔ G`. Let *c* be the nucleotide to be complemented. Then the following is true 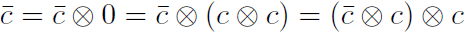, but also 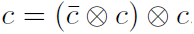. So *c* can be complemented back and forth with the same operation. Furthermore, 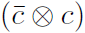 simplifies to a constant value, reducing the complement to one machine instruction.

So two magic constants suffice to complement all four nucleotides: `21 = (A ⊗ T)` and `4 = (C ⊗ G)`. To pick the right constant for complementing, a trick similar that is in the hashing section above, can be used; `C` and `G` have their bit 2 set, whereas `A` and `T` do not.

~~~
**char** *\*revcomp*(**const char** * *forward*, **char** * *reverse*, **size_t** *len*)
{
    **for (size_t** *k* = 0; *k < len; k*++) *{*
        **char** *c* = *forward*[*k*];
        **char** *magic* = *c* & 2 ? 4 : 21;

        *reverse* [*len – k –* 1] = *c*^ *magic;*
    }

    *reverse*[*len*] *=* ’\0’;
    **return** *reverse;*
**}**
~~~

This code is surprisingly short and could be compacted even further. It contains only one branch and very simple instructions, making it fast and vectorizing-friendly.

### 2.5 Mutations and Transversions

As a final task we focus our attention on comparing genomes, specifically, counting and classifying mutations. Given two genomes *s,q* ∈ Σ*, with equal length, we are interested in the number of transversions separating the two sequences.

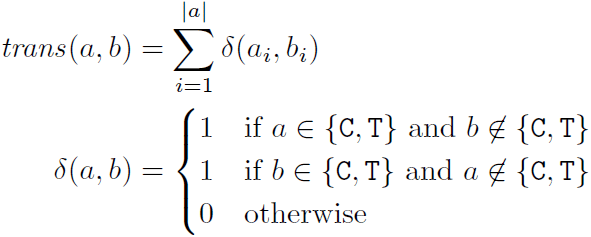

To account for a transversion, exactly one of the two bases has to be a pyrimidine, and the other purine. A character *c* is a pyrimidine in ASCII if *c* + 1 has a 1 at bit 3. For purines that bit has the value 0. So incrementing the characters by one allows for easy characterization of bases. With a bitwise-exclusive-or we can check if only one of the two bases is a pyrimidine.

~~~
**double** *transversions*(**const char** \**subject*, **size_ t** *length*, **const char** * *query*)
{
    **Size_ t** *transversions* = 0;

    **for** (**size_t** *k* = 0; *k < length; k*++) {
        if (((*subject*[*k*] + 1) ^ (*query*[*k*] + 1)) & 4) {
            *transversions*++;
        }
    }

    **return** (**double**)*transversions* / *length*;
}
~~~

## 3 Results

To evaluate the performance of the given methods, we simulated sequences with 100,000 nucleotides. On each sequence a method is run often enough to gain statistical confidence using the benchmark library by Google [3]. This process is repeated a number of runs, from which the *minimum* is chosen as the best run-time[1].

Also, we inspect the runtime characteristics of different methods using the perf tools [2]. These allow measuring the instructions per cycle, branches and other features of a program.

### 3.1 GC-Content

In Section 2.2 we describe, how the GC-content of a sequence can be computed with fewer comparisons than necessary at first glance. We created two methods, one representing the simple way of counting and the other the explicit twiddling way. As modern compilers optimize the generated code, we do not expect any significant difference between the two methods. As a third method we use a table look up.

Figure 1 shows the benchmarking results for the three methods. Using the GNU Compiler all methods are almost equally fast. However, Clang fails to optimize the simple method. It still makes one extra comparison leading to many more branches and thus a high number of branch misses. This leads to a roughly 25 times slower function.

**Figure 1.**
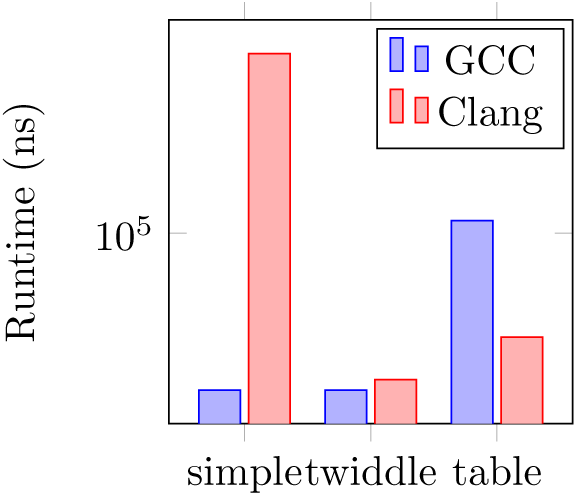
Computing the GC-Content. Runtimes of different methods to determine the GC-content from from sequences with 100,000 nucleotides. Todo: fix the y-axis.

### 3.2 Hashing

We compare the two hashing procedures defined in Section 2.3, with one using a look-up table. One uses a big switch statement to map the nucleotides to an index. The other exploits the ASCII representation of the characters to achieve the same result, much faster. We measured the performance by having both methods compute the hash of all *k*-mers in a long sequence with *k* = 16.

As we would have hoped, the twiddling method is faster than the simple switch, by a factor of 3. This is due to different reasons. The twiddle method utilizes more instruction-level parallelism. On an Intel Core i5-5200U the simple method executes about 1.19 instructions per cycle. However, using twiddling, that number can be ramped up to 3.75. On the contrary, the number of branches and especially the number of branch misses goes down. Whereas for the simple method 25% of all branches are missed, it is only 0.02% for twiddle.

Using a look-up table is even faster than twiddling by roughly 20%. This is mainly due to a greatly reduced number of instructions.

**Figure 2.**
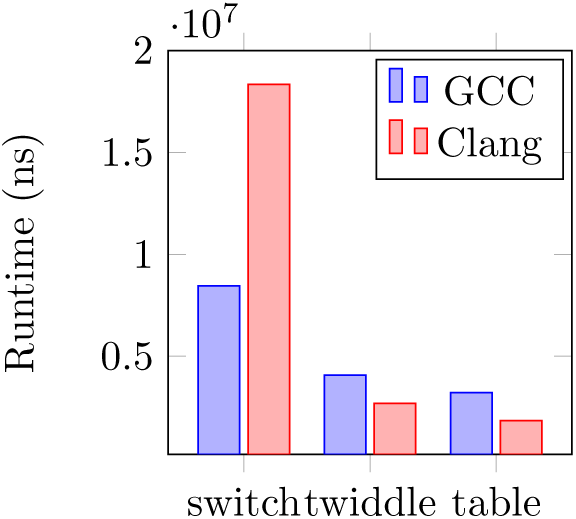
Runtime of different k-mer hashing procedures.

### 3.3 Reverse Complement

Here we compare four different ways of computing the reverse complement. As a baseline, again we use our simple switch-statement. twiddle is the method using xors as presented in Section 2.4. A third method used by programs such as Blast and MUMmer uses a table lookup. Lastly, one method (two step) splits the process into two parts. First, all nucleotides are complemented, then the string is reversed.

It can be seen in Figure 3 that the switch method is by far the slowest. All other methods are at least one order of magnitude faster. Among them the table lookup is the slowest, followed by a two-step process. Twiddling leads to the best performance. All results are the same across compilers.

**Figure 3.**
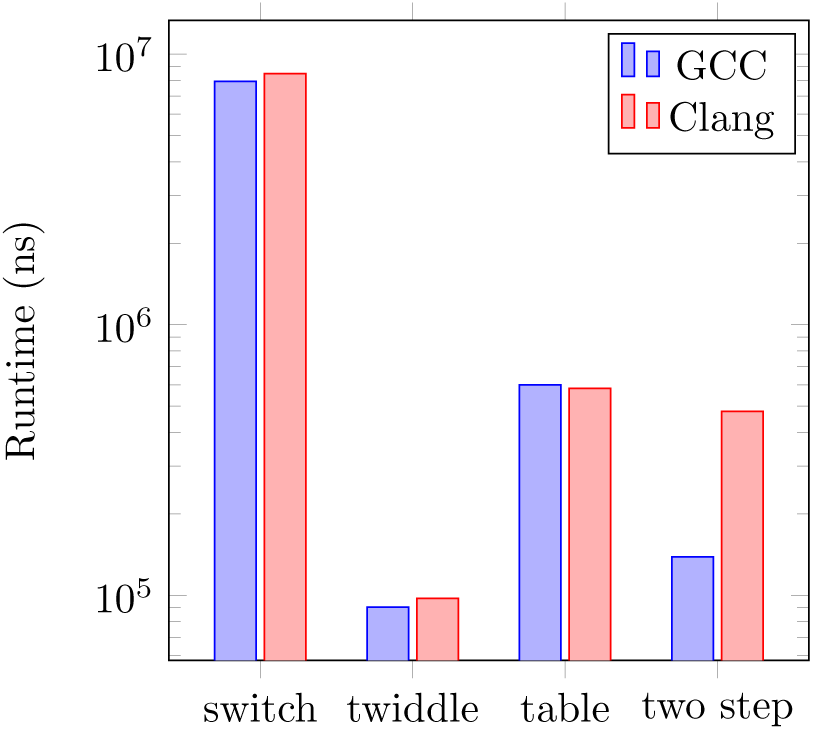
Reverse Complement. Runtime of di_erent methods for computing the reverse complement of a sequence with 100,000 nucleotides.

### 3.4 Transversions

To compare the performance of different ways of counting transversions, we simply choose counting mutations as a baseline.

The simple way of counting transversions cannot be vectorized and thus is more than ten times slower than counting only mutations. However, the new twiddling method presented in Section 2.5 can be vectorized, an thus is almost as fast as the first method. Likewise, using a look-up table cannot be vectorized as thus is slower than twiddling, but ten times faster than a simple comparison.

**Figure 4.**
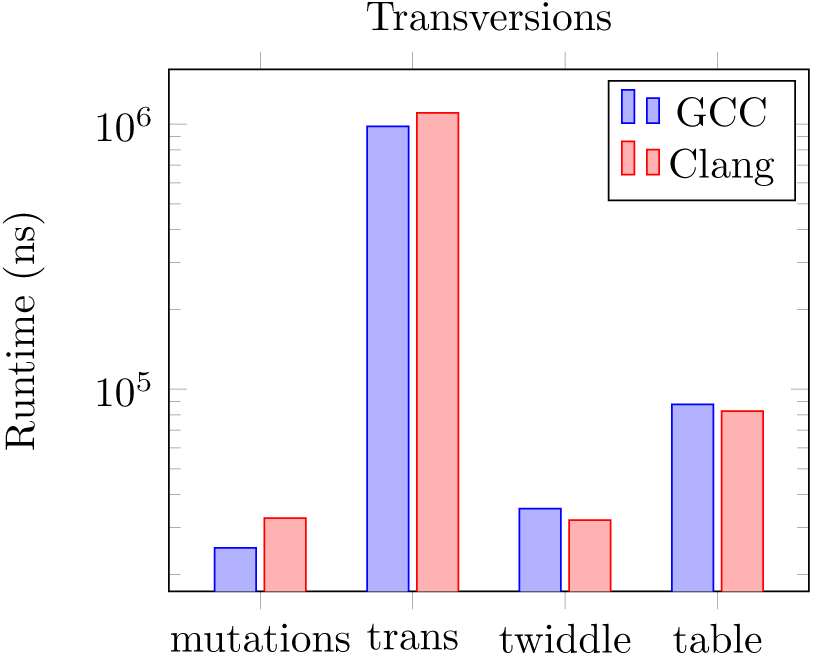
Counting Transversions. Runtime of di_erent methods for counting mutations and transversions. Shown is the minimum of 3 runs.

## References

[1] A. Alexandrescu. Writing fast code i, 2015.

[2] L. K. Developers. perf. Technical report, 2016.

[3] Google. A microbenchmark support library. Technical report, GitHub, 2016.

[4] J. Longinotto. up2bit format, 2015.

